# COVID-19 and the abrupt shift to remote learning: Impact on grades and perceived learning for undergraduate biology students

**DOI:** 10.1101/2021.03.29.437480

**Authors:** K. Supriya, Chris Mead, Ariel D. Anbar, Joshua L. Caulkins, James P. Collins, Katelyn M. Cooper, Paul C. LePore, Tiffany Lewis, Amy Pate, Rachel A. Scott, Sara E. Brownell

## Abstract

Institutions across the world transitioned abruptly to remote learning in 2020 due to the COVID-19 pandemic. This rapid transition to remote learning has generally been predicted to negatively affect students, particularly those marginalized due to their race, socioeconomic class, or gender identity. In this study, we examined the impact of this transition in the Spring 2020 semester on the grades of students enrolled in the in-person biology program at a large university in Southwestern United States as compared to the grades earned by students in the fully online biology program at the same institution. We also surveyed in-person instructors to understand changes in assessment practices as a result of the transition to remote learning during the pandemic. Finally, we surveyed students in the in-person program to learn about their perceptions of the impacts of this transition. We found that both online and in-person students received a similar small increase in grades in Spring 2020 compared to Spring 2018 and 2019. We also found no evidence of disproportionately negative impacts on grades received by students marginalized due to their race, socioeconomic class, or gender in either modality. Focusing on in-person courses, we documented that instructors made changes to their courses when they transitioned to remote learning, which may have offset some of the potential negative impacts on course grades. However, despite receiving higher grades, in-person students reported negative impacts on their learning, interactions with peers and instructors, feeling part of the campus community, and career preparation. Women reported a more negative impact on their learning and career preparation compared to men. This work provides insights into students’ perceptions of how they were disadvantaged as a result of the transition to remote instruction and illuminates potential actions that instructors can take to create more inclusive education moving forward.

## Introduction

In the early months of 2020, the COVID-19 pandemic led to an unprecedented disruption of the normal mode of course instruction across most institutions of higher education. In the United States, most universities abruptly stopped conducting in-person classes and closed their campuses in March 2020 [1,2]. Mid-semester, many students and instructors were forced into learning and teaching remotely, respectively, for the first time due to the need for social distancing as a response to the pandemic [3,4]. Syllabi, teaching approaches, and assessments had to be modified to account for this altered mode of learning; most instructors only had one to two weeks to redesign their courses before remote instruction began. This abrupt shift to remote learning has been distinguished from online learning in general [5] and it is commonly assumed that this abrupt shift adversely affected student learning [6,7]. There are many factors directly associated with the shift to remote learning that could have affected student learning [5,7], which are in addition to the stress experienced by students in other aspects of their lives affected by the pandemic (e.g., health, employment, isolation, issues of inequality).

The pandemic affected people across various social identities such as age, nationality, racial/ethnic background, LGBTQ+ status, and socio-economic status. Despite being termed as “the great equalizer” by politicians like New York’s Governor Andrew Cuomo and celebrities such as Madonna [8,9], it had differential impacts on people along the lines of power and privilege in our society due to various systems of oppression including, but not limited to, racism, classism, sexism, and ableism [9–13]. In the United States, case and death rates have been higher among Black, Hispanic/Latinx, and Native American people than white people [14–16]. COVID-19 infections and deaths were also higher for people living in areas with higher poverty levels compared to areas with little or no poverty [17,18]. Further, these more vulnerable communities experienced more negative financial impacts such as job losses or reduced working hours due to the economic shutdowns [19]. Moreover, some studies have reported more negative mental health impacts of the pandemic on women, Hispanic, and Asian people [20], and on people living in lower-income households [21]. When considering the educational impact of this crisis, it is important to ask if these differential medical and financial impacts contributed to more negative educational consequences for students with marginalized social identities.

In addition to health and financial impacts, several other factors may have differentially exacerbated the negative effects of the COVID-19 pandemic on student learning in Spring 2020. Losing access to student housing and meal plans contributed to housing and food insecurities for many students, including low-income students and international students [22,23], and heightened housing and food insecurities impacted off-campus students as well [24]. Moreover, poor internet connection and lack of a quiet or safe space to study made it more difficult for students to complete their assignments and succeed during remote instruction [25–28]. For example, one recent study of college students in introductory sociology courses showed that more than 50% of all students experienced occasional internet problems during remote learning in Spring 2020 [29]. In the same study, about 90% of the students reported distractions in their new workspace and about 65% of the students reported the lack of a dedicated workspace [29]. While these issues negatively affect all students, students from low-income families are disproportionately impacted by poor internet connections or distracting environments. Another factor that likely affected remote learning in Spring 2020 is additional caregiving responsibilities necessitated by remote learning in K-12 schools and greater health risks for older family members [30]. These additional responsibilities would reduce time for coursework and could affect academic outcomes. Likely due to societal gender roles that assume women take on primary caregiving, these responsibilities are reported to have disproportionately affected women [30–32]. The privilege of staying at home or having safe working conditions to reduce the risk of exposure to COVID-19 has also been shaped by axes of power in our society [33–35]. Needing to work jobs that require frequent interaction with others at places such as grocery stores and pharmacies is yet another element influencing student learning during the pandemic, especially for Black, Hispanic/Latinx, immigrant students, and those from low-income households [36]. Working such jobs could increase students’ risk of exposure to the virus and may cause greater anxiety in their daily lives [37,38]. All these factors are likely to differentially affect students depending on their locations along the various axes of power and privilege.

A limited number of studies have examined the educational impact of the pandemic on students. Several publications have reported that students were less engaged [39] and struggled with their motivation to study after the transition to remote learning in Spring 2020 [25,29,40,41]. One study on public health students at Georgia State University did not report lower motivation among students [42], perhaps because of the heightened awareness of the relevance of public health during a global pandemic. It has also been demonstrated that the transition to remote learning had a negative impact on student relationship-building, specifically the extent to which students interact with each other in and out of class [25,43], and on students’ sense of belonging in the class [25]. In response to the pandemic, several universities changed course policies to extend the deadline for course withdrawals or to allow greater access to pass/fail grading options [44]. Villanueva and colleagues [28] found higher course withdrawal rates among general chemistry undergraduates after students were offered an extended deadline for withdrawing from the course. Despite these negative student experiences, some studies have reported small increases in student grades in Spring 2020 compared to similar courses in previous years [45–47].

There is some evidence for differential impacts of the transition to remote learning for students with different social identities. For example, a report based on survey data from 600 undergraduates in STEM courses across the US showed that women, Hispanic students, and students from low-income households experienced major challenges to continuing with remote learning more often than men, white students, and students from middle- or high-income households, respectively [25]. Another survey study found that the likelihood of lower-income students delaying graduation because of COVID-19 was 55% higher than higher-income students [48]. Additionally, Gillis and Krull [29] reported that women experienced challenges such as lack of a dedicated workspace more often than men, while non-white students experienced anxiety over personal finances and access to medical care more often than white students.

In contrast to students in in-person degree programs whose mode of learning changed drastically, the crisis did not fundamentally change the mode of learning for students who were already enrolled in fully online degree programs. Although other aspects of the lives of online students were still affected by the pandemic, online learning was not new to them or their instructors, courses did not need to be modified halfway through the term, and students expected to complete all coursework remotely when they signed up for the course. Therefore, comparing the impact of the pandemic on the grades of online and in-person students might allow us to tease apart the influence of the rapid transition to online learning from the stress of living through a global pandemic. One prediction would be that online students would experience less of a negative impact on learning due to the pandemic compared to their in-person counterparts because their educational modality did not change. An alternative prediction is that the differences in the student populations online and in-person, specifically the higher percentage of individuals in the online program who hold one or more marginalized social identities and may be more vulnerable to the negative effects of the pandemic outside the class, would lead to greater negative impacts for online students as a result of the COVID-19 pandemic. Specifically, we know that the percentage of women, older students, students who are primary caregivers, and students from low-income households are consistently higher in online programs compared to in-person programs [49–51]. These are groups that have been unequally disadvantaged during the pandemic in general. Therefore, it is important to control for demographic variables when comparing the effects of the COVID-19 pandemic on grades between students in online and in-person degree programs.

## Current Study

The biology program at Arizona State University (ASU) offers a unique opportunity to examine the impact of the emergency transition to remote learning on undergraduates. First, ASU offers equivalent in-person and fully online biology degree programs that have aligned curricula. This allows for comparison of the experiences of students in an in-person program transitioned to remote learning, to the experiences of students enrolled in an online program prior to the COVID-19 pandemic. In this study, following the recommendation from Hodges et al. [5] we use the term “remote” to refer to in-person courses that transitioned abruptly to online instruction, while using the term “online” for courses that were designed to be online from the beginning. One important difference between the online and in-person programs after the transition to remote learning in Spring 2020 was that courses in the online program were fully asynchronous. In contrast, the courses in the in-person program were generally taught synchronously using web conferencing (e.g., Zoom) for lectures and typical in-class activities.

Second, ASU has a large, diverse population of students that allows for the examination of the extent to which the transition affected students with different social identities. Science, technology, engineering, and math (STEM) disciplines, such as biology, have long been exclusionary spaces dominated by relatively wealthy white men [52–54]. Underrepresentation of women, people of color, people with disabilities, and people with low socioeconomic status is well documented in the sciences [55]. Therefore, it is important to examine the impact of the transition to remote learning on STEM students with social identities historically underrepresented in the sciences, for which ASU’s biology program provides a suitable context.

This study uses course grades during the Spring 2020, Spring 2019, and Spring 2018 semesters and survey data from instructors and students about the Spring 2020 semester to examine the impacts of the abrupt transition to remote learning due to COVID-19 during the Spring 2020 semester.

Specifically, our research questions were:

1. Did the abrupt transition to remote learning due to the COVID-19 pandemic affect grades for undergraduate students in an in-person biology program during the Spring 2020 semester? Was this effect on grades different from that found in the equivalent online biology program during Spring 2020? To what extent did the abrupt transition to remote learning disproportionately affect students with identities historically underrepresented in STEM?

2. What changes did in-person biology instructors make to their assessment practices after the abrupt transition to remote learning in Spring 2020 and to what extent do these explain any differences in student grades observed?

3. To what extent do in-person biology students perceive that their learning, interactions with peers and instructors, career preparation, interest in science, and feeling a part of the biology community were affected because of the abrupt transition to remote learning? To what extent did the abrupt transition to remote learning disproportionately affect these perceptions for students with identities historically underrepresented in STEM?

## Positionality of the authors

We acknowledge that our own identities influence the research questions that we ask and how we may interpret the data. Our author team includes individuals who identify as men, women, white, South Asian, Jewish, first-generation college-goers, first-generation immigrants, and members of the LGBTQ+ community; members of our team grew up in middle class families in the United States, except KS who grew up in India. All the authors are committed to diversity, equity and inclusion in the sciences and conduct education research focused on equity. This paper was motivated by our concerns regarding social inequities and how they are perpetuated and, in some cases, amplified in undergraduate science classrooms.

## Methods and Results

### Research Question 1: Assessing the impact of the abrupt transition to remote learning due to the COVID-19 pandemic on grades for undergraduate students in an in-person biology program compared to an online biology program

#### Research Question 1, Methods

To study the impact on student course grades that resulted from the shift to remote learning during the COVID-19-impacted Spring 2020 semester, we obtained course grades from the university registrar for Spring 2020 and compared these grades to two spring semesters prior to the pandemic: Spring 2019 and Spring 2018. The population of interest is undergraduate biology majors enrolled in either the in-person biology degree program or the fully online biology degree program. Therefore, we obtained course grades for 42 STEM courses that are commonly taken by students in these biology majors, including general biology courses, biochemistry, chemistry, physics, mathematics, and statistics. See Table S1 for the full list of courses.

Our grades analysis included a total of 25,100 student-course enrollments, with 8,323 from the Spring 2020 pandemic semester and the remainder from Spring 2018 or 2019. Of these, 19,181 course enrollments were in-person courses and the remaining 5,919 were online degree program courses. Course grades were analyzed on a 0–4.33 scale (A+ = 4.33, A = 4.0, A- = 3.66,…, E = 0). Grades other than A–E were excluded from analysis; this was a total of 2,404 student-course enrollments, or 9.6% of the total dataset. In Spring 2018 and 2019, these excluded grades are almost exclusively W or “withdraw” grades. In response to the unique circumstances of the pandemic, some instructors assigned the “Y” grade which indicates “Satisfactory” work at a level of C or higher. In Spring 2020, about a third of the non-letter grades were Y grades. The combined proportion of non-letter grades held steady in Spring 2020 compared to 2018 and 2019 in online courses and increased slightly in Spring 2020 for in-person courses. The withdrawal percentage declined, and the Y percentage rose both online and in-person. We cannot say definitively how many of the students who received a Y grade would have chosen to withdraw if this option had not been available. The decision to remove these grades from analysis is consistent with prior studies [51,56]. To control for prior academic performance, we use “GPAO,” which refers to a student’s grade point average in other courses, including both STEM and non-STEM courses [56,57].

We obtained student demographic information from the registrar in addition to course grades (summarized in Table 1). The categories of interest for this study are gender, race/ethnicity, and two proxies for socioeconomic status (college generation status and federal Pell grant eligibility). The transition away from an in-person lecture and having to adapt to a large change mid-semester could also have negatively affected the learning of students with disabilities [7] as changing learning environments have presented novel challenges for deaf and hard of hearing students [58] and students with disabilities more broadly [7]. However, because we are using institutional data in these analyses and data on disabilities is protected by federal law, we were not able to examine the impact of the transition on students with disabilities in this study, nor were we able to explore other identities not routinely collected by the university registrar.

**Table 1.** Demographics for students in the in-person and online course grades data set. Pell eligibility and college generation status are included as proxies for socioeconomic status. BLNP refers to Black, Latinx, Native American, and Pacific Islanders.

|  | In-Person Students | Online Students |
| --- | --- | --- |
| Characteristic | N = 4,671<sup>1</sup> | N = 2,586<sup>1</sup> |
| Gender |  |  |
| Man | 1,652 (35%) | 675 (26%) |
| Woman | 3,019 (65%) | 1,910 (74%) |
| Other | 0 (0%) | 1 (<0.1%) |
| BLNP |  |  |
| N | 3,058 (65%) | 1,662 (64%) |
| Y | 1,613 (35%) | 924 (36%) |
| Race/Ethnicity |  |  |
| White | 2,242 (48%) | 1,431 (55%) |
| Asian | 625 (13%) | 128 (4.9%) |
| Black | 245 (5.2%) | 224 (8.7%) |
| Characteristic | N = 4,671<sup>1</sup> | N = 2,586<sup>1</sup> |
| Hispanic | 1,168 (25%) | 547 (21%) |
| Native | 65 (1.4%) | 39 (1.5%) |
| Two or more races | 273 (5.8%) | 159 (6.1%) |
| Decline to state | 53 (1.1%) | 58 (2.2%) |
| Pell Eligible |  |  |
| N | 2,689 (58%) | 1,079 (42%) |
| Y | 1,982 (42%) | 1,507 (58%) |
| College Generation Status |  |  |
| Continuing Generation | 3,169 (68%) | 1,458 (56%) |
| First Generation | 1,502 (32%) | 1,128 (44%) |
| <sup>1</sup>n (%) |  |  |

To determine the direction and significance of the effect of the shift to remote learning on student grades, we performed a linear mixed-effects regression on the numerical course grades. The fixed effects in the model included a dummy variable for the Spring 2020 (“COVID-19”) semester, whether the student was enrolled in the in-person or online degree program, an interaction between these two variables, and the GPAO term. We included random effect terms for course section and student. These terms provided modest improvement to the models with a combined intraclass correlation coefficient equal to 0.256.

To determine the direction and significance of the effect of the shift to remote learning on grades received by students with identities historically underrepresented in STEM, we added interaction terms between the dummy variable for the Spring 2020 (“COVID-19”) semester and each of the demographic terms to the model described above. We again controlled for GPAO and included random effect terms for course section and student in this model (see Table S2 for model specifications).

#### Research Question 1, Results

Overall, our linear mixed effects regression results show that the Spring 2020 semester was associated with a positive grade shift of 0.41 grade units. Students earned higher grades in Spring 2020 courses compared to students enrolled in those courses in Spring 2019 and Spring 2018. Results also show that this Spring 2020 grade effect was not significantly different between the online and in-person programs (Table 2). The online program is also associated with lower course grades overall, which is consistent with our prior work [51].

**Table 2.** Linear mixed effects regression results showing Spring 2020 (“COVID-19”) effect and its interaction with instruction mode.

| Variable | Beta | 95% CI<sup>1</sup> | p-value |
| --- | --- | --- | --- |
| (Intercept) | 0.12 | 0.05, 0.20 | 0.002 |
| GPAO | 0.89 | 0.87, 0.91 | <0.001 |
| Spring 2020 | 0.41 | 0.36, 0.46 | <0.001 |
| Campus |  |  |  |
| In-Person | — | — |  |
| Online | -0.27 | -0.38, -0.15 | <0.001 |
| Spring 2020 * Campus |  |  |  |
| Spring 2020 * Online | -0.11 | -0.33, 0.11 | 0.3 |

<sup>1</sup>CI = Confidence Interval

Our regression model testing for the presence of negative interaction effects between demographic groups and the Spring 2020 semester showed no significant negative interactions for any of the demographic variables that we examined, including gender, race/ethnicity, and socioeconomic status (Table S3). Contrary to our prediction, the model shows positive, but mostly non-significant, interaction effects for all groups compared to their historically overrepresented counterparts. The two statistically significant interactions showed women to have a Spring 2020 effect 0.05 greater than men and Pell-eligible students to have an effect 0.08 greater than non-Pell-eligible students.

### Research Question 2: Understanding biology instructor changes to assessment practices in the Spring 2020 semester when they transitioned to remote learning and their effects on differences in student grades

#### Research Question 2, Methods

To better understand why the COVID-19 pandemic did not negatively affect course grades in the Spring 2020 semester for students who had to transition to remote learning, we sought to understand what steps in-person biology instructors took to ensure that their students could achieve the course goals after the abrupt transition to remote learning. To explore this, we created a survey with several open-ended questions regarding changes in instructional practices, such as modes of interaction with students and assessments used after the transition to remote learning (a copy of the survey questions analyzed is provided in the Supplemental Materials). Of the 132 biology instructors recruited to participate, 27 instructors responded to the survey (20% response rate). Faculty members were recruited first via email, and verbally encouraged to participate at several follow-up virtual events attended by many of those in the recruitment group.

Building on the open-ended responses from the first instructor survey, we created a second survey that asked in more detail about instructional changes in response to the pandemic. To assess cognitive validity, we conducted two think-aloud interviews with biology faculty members who taught in person during Spring 2020 and had to transition to remote learning [59]. These think-aloud interviews indicated that the instructors understood the questions. We then distributed this revised survey to all biology instructors who taught in-person courses in Spring 2020 (n=132). In the event that they taught multiple courses, the survey asked them to respond based on their largest course size. The survey first asked instructors to identify any changes they made in their course. This question used a multiple-selection format with a) 24 options provided, b) an option to say that no changes were made, and c) an option to describe other changes not listed. The survey also asked instructors to report the extent to which they tried to reduce cheating in their course, the extent to which they made their course more flexible, and the extent to which they made their course easier. Each of these questions was answered using a six-point Likert scale from strong agreement to strong disagreement with no neutral option and they were asked to explain each answer (a copy of the survey questions analyzed is provided in the Supplemental Materials). While instructors also experienced many of the same personal challenges resulting from the pandemic that students did, our focus was on the student experience and therefore we only asked instructors about instructional changes.

A total of 43 out of the 132 biology instructors who were contacted completed the second survey (33% response rate) based on their experiences teaching an in-person biology course that shifted to fully remote instruction in the Spring 2020 semester. Of these, 18 had taught an in-person course that transitioned during Spring 2020 with at least 100 students. Our analysis will focus on these large courses because these instructors are subject to greater practical constraints when considering how to shift instruction to remote learning and because the larger sizes mean that a greater number of students in total are impacted by these decisions.

To understand the extent to which changes in assessment practices made by instructors might explain differences in student grades in Spring 2020 compared to previous semesters, we examined data from 10 instructors who responded to our survey who had taught the same course in Spring 2020 and either Spring 2019 or 2018. We performed course-level linear regressions on the relative grade difference using the following variables as predictors: total number of changes made, use of lockdown browsers for exams, whether they made efforts to reduce cheating, and whether they worked to make the course easier. All variables were dichotomous except number of changes made. The question about making the course more flexible was not included because all ten of the instructors who had taught the same course in Spring 2020 and Spring 2019 or 2018 agreed with this question.

#### Research Question 2, Results

Overall, most instructors reported making changes to their in-person courses when they needed to transition to remote learning during the COVID-19 pandemic, including being more flexible and making the course easier (Table 3). Focusing on the large courses, about 60% of instructors agreed that they took steps to reduce cheating. Nearly all large course instructors (94%) agreed that they made changes to be more flexible to help students who were experiencing challenges and most (78%) agreed that they made it easier for students to do well.

**Table 3.** Summary of in-person instructor survey responses about the changes they made to their course after the transition to remote learning in Spring 2020.

| Survey Item | Large Class<br>Instructors<br>N = 18<sup>1</sup> | Small Class<br>Instructors<br>N = 25<sup>1</sup> | All Instructors<br>N = 43<sup>1</sup> |
| --- | --- | --- | --- |
| Made more flexible | 94% | 84% | 88% |
| Made easier | 78% | 56% | 65% |
| Tried to reduce cheating | 61% | 32% | 44% |
| Number of changes selected | 4.7 (3.2) | 3.1 (3.5) | 3.8 (3.4) |
| Zero changes selected | 5.6% | 24% | 16% |

<sup>1</sup>%; Mean (SD)

Instructors were also asked to select the changes they made to their largest in-person course that had to transition to remote learning in that semester from a list of 24 options (Table 4, See Supplemental Table 4 for the full set of options). On average, instructors of large courses selected about five changes. The most frequently selected changes were generally related to time and deadline extensions as well as conducting open-book exams. Changing the weighting or number of exams or changing the difficulty of questions on quizzes or exams were less commonly selected. Thirteen respondents added open-ended comments in addition to the provided choices. Five of these related to changes needed to replace planned fieldwork or labs. The remainder detailed specific content-related adjustments or discussed changes to increase instructor availability to students.

**Table 4.** Frequencies of selection of fixed choice options for course changes by in-person instructors after the transition to remote learning in Spring 2020. This table only shows options chosen by ≥25% of respondents; for full results, see Table S4).

| Response Option | Frequency<br>N = 43 |
| --- | --- |
| Gave individual students extensions on deadlines for out-of-class assignments that I wouldn't have normally provided | 37% |
| Extended the deadline or allotted more time than I usually provide to complete out-of-class assignments | 33% |
| Increased the amount of time students were allotted to complete a quiz or exam | 33% |
| Gave students more opportunities to miss class and not lose participation/attendance points but still gave participation/attendance points for class | 26% |
| Reduced or eliminated penalties for out-of-class assignments that were submitted late | 26% |
| Changed assessments such as exams or quizzes from closed-book to open-book | 26% |
| In addition to delivering my content online I made a significant change to my course that is not reflected above | 30% |

Within our subset of surveyed instructors who had taught the same class in Spring 2020 and one of the prior Spring terms, our linear regression showed that none of the instructional changes in assessment practices were significant predictors of the difference in grades received by the analyzed students in Spring 2020 compared to previous two Spring semesters. Greater instructor flexibility could be associated with the increase in grades across all courses, but we were not able to test this relationship because all ten of the instructors in this subset reported increasing flexibility in their courses.

### Research Question #3: Understanding the impact of the abrupt transition to remote learning due to COVID-19 on biology student perceptions of learning, interactions with peers and instructors, career preparation, interest in science and feeling a part of the biology community

#### Research Question 3, Methods

Although the transition to remote learning for students who were in the in-person biology degree program in Spring 2020 did not have adverse effects on their grades, likely in part because instructors made changes to their courses, we wanted to explore student perceptions of learning during Spring 2020. To do so, we surveyed students during Fall 2020 to ask specifically about their experiences during the Spring 2020 semester when their in-person courses rapidly transitioned to remote learning.

##### Student survey development

To assess the perceptions of biology majors who experienced the rapid transition from in-person to remote instruction in Spring 2020, we developed a survey that contained both closed-ended and open-ended questions. We asked students to think about the largest biology course they took in the Spring 2020 semester to answer the survey questions that were course-specific, (i.e., impact on grades, impact on learning, and perceived instructional changes). For the rest of the survey questions, students were asked to think about all the in-person biology courses they took in Spring 2020. To assess cognitive validity of survey items, we conducted six think-aloud interviews with undergraduate students and iteratively revised survey items until no further changes were suggested [59]. The final survey contained questions about the perceived impact of the rapid transition to remote learning on student learning, grades, interest in their biology major, interest in learning about scientific topics, feeling a part of the biology community at the university, and career preparation. Each question was answered using a seven-point scale from “strong negative impact” to “strong positive impact.” In addition, we asked about the impact of the transition on the amount of time spent interacting with instructors and other students, and the amount of spent time studying. These items were also answered using a seven-point scale ranging from “greatly decreased” to “greatly increased.” During our think-aloud interviews with undergraduate students, the necessity of a “neutral” option for these survey items was brought up by multiple students. Therefore, we used a seven-point scale for these items instead of the six-point scale used in our instructor survey. We also asked students about perceived instructional changes to the course in terms of measures to prevent cheating, increase flexibility, and make the course easier. These were on a six-point scale from “strongly agree” to “strongly disagree” with no neutral option for consistency with the instructor survey (see Supplemental Materials for the analyzed survey questions).

We included some demographic questions at the end of the survey so we could test for any differential effects on student experience by social identities, specifically gender, race/ethnicity, college generation status and eligibility for federal Pell grants. For race/ethnicity, we asked students two questions: whether they identified as Hispanic/Latinx and whether they identified as Black/African American, Native American/Alaska Native, or Native Hawaiian/Pacific Islander. Students that selected “yes” to either of these questions were grouped together as BLNP for our analyses. We grouped students in this manner because all these groups are historically underrepresented in the sciences and our sample sizes for the student survey were not large enough to allow us to disaggregate race/ethnicity data.

##### Student survey distribution

In Fall 2020, we used a convenience sampling approach to recruit eight biology instructors who agreed to distribute our survey to students in their classes. The survey was sent to a total of 1,540 students in these eight courses and students were offered a small amount of extra credit for completing the survey. A total of 798 students completed the survey, resulting in a response rate of 51.8%. However, only 601 of these students were enrolled in the in-person biology degree program in Spring 2020. Of these students, 70 reported that they did not take any biology courses in Spring 2020, so they were not included in any course-specific analyses. After removing these students and removing 21 students with missing data, we were left with responses from 510 students who had taken in-person biology courses that had transitioned to remote learning in Spring 2020 that we used for our analyses (Table 5). Students were asked to think about the largest in-person biology course they took in Spring 2020 for the survey. This gave us data about student experiences in 25 Spring 2020 courses, although for 13 of these Spring 2020 courses, we had fewer than 10 respondents.

**Table 5.** Demographics for student survey respondents. Three of the women among the survey respondents also identified as non-binary. One of the men also identified as non-binary and as transgender. BLNP refers to Black, Latinx, Native American, and Pacific Islanders. Pell eligibility and college generation status are included as proxies for socioeconomic status.

|  | Student survey respondents |
| --- | --- |
| Characteristic | N = 510<sup>1</sup> |
| Gender |  |
| Man | 165 (32%) |
| Woman | 345 (68%) |
| Other | 0 (0%) |
| BLNP |  |
| N | 358 (70%) |
| Y | 152 (30%) |

|  | Student survey<br>respondents |
| --- | --- |
| Race/Ethnicity |  |
| Asian | 98 (19%) |
| Black | 17 (3.3%) |
| Hispanic | 108 (21%) |
| Native | 6 (1.2%) |
| Two or more races | 38 (7.5%) |
| White | 219 (43%) |
| Decline to state | 24 (4.7%) |
| Pell Eligible |  |
| N | 336 (66%) |
| Y | 174 (34%) |
| College Generation<br>Status |  |
| Continuing<br>Generation | 357 (70%) |
| First Generation | 153 (30%) |
| <sup>1</sup>n (%) |  |

##### Student survey analyses

We calculated the total percentage of students that reported negative impacts on their learning, amount of time studying and interacting with peers and instructors, career preparation, interest in science and feeling a part of the biology community. To analyze the open-ended data, we used open-ended coding methods to identify themes that emerged from student responses [60]. We used constant comparison methods to develop the coding scheme; student responses were assigned to a category and were compared to ensure that the description of the category was representative of that response and not different enough to require a different category. Inter-rater reliability was established by having two coders (S.E.B. and R.A.S.) analyze 20% of the data, after which one person coded the rest of the data. For student perceptions of the positive impact of the transition to remote instruction on learning codes: Two raters compared their codes and their inter-rater reliability was at an acceptable level (k = 0.88). For student perceptions of the negative impact of the transition to remote instruction on learning codes: Two raters compared their codes and their inter-rater reliability was at an acceptable level (k = 0.88). We report out any code that at least 10 students mentioned.

For eight of the Spring 2020 courses in our dataset, we had data from both the instructor and more than 10 students for each course. For these courses, we assessed if student responses to perceived instructional changes to the course aligned with the instructional changes as reported by the instructors. We analyzed the strength of this relationship through Pearson product-moment correlations between the percent of students agreeing with each statement and the strength of the instructor’s agreement using a Likert scale.

To examine demographic differences in the perceived impact on students, we used ordinal mixed model regressions with the Likert scale option chosen by students as the outcome and gender, race/ethnicity, Pell-eligibility, and first-generation to college status as predictors. We used course section as a random effect with varying intercepts in all the models to account for the nested nature of our data. We used the R regression package *ordinal* [61] for these analyses.

#### Research Question 3, Results

About 56% of students reported that they think the transition to remote learning negatively impacted their grade, even though our grade analysis did not indicate that this was likely.

However, almost 70% of students said that transition to remote learning negatively impacted their learning in the same course (Fig 1). We analyzed the reasons why students felt that the transition to remote instruction either positively or negatively affected their learning (Tables 6 and 7). For the 30% of students who thought that it positively impacted their learning, they said it did so because lectures were recorded so they could review them or see more of them (18.3%), they felt as though they could learn at their own pace (15.0%), they felt like remote learning allowed them to engage with the material in a more active learning way (11.7%), or they felt more comfortable learning at home as opposed to in a large classroom (8.3%). There was also a subset of students who felt as though they had more time in general during the pandemic, which allowed them to focus more on studying (16.7%). For the 70% of students who reported that the pandemic negatively impacted their learning, 27.2% of students reported that they felt as though they understood less and remembered less during remote instruction (Table 7). Students also reported a loss of concentration or focus (26.6), fewer opportunities to interact with others and ask questions (17.0%), and having less motivation or interest (9.9%). Less common responses included: feeling overwhelmed by greater amounts of work after the transition to remote learning (5.9%), lack of hands-on learning, particularly in lab courses (4.0%), general stress associated with the pandemic that increased distractions outside of coursework (3.7%), procrastination and less accountability (3.1%), and technical issues (2.8%).

**Fig 1.**
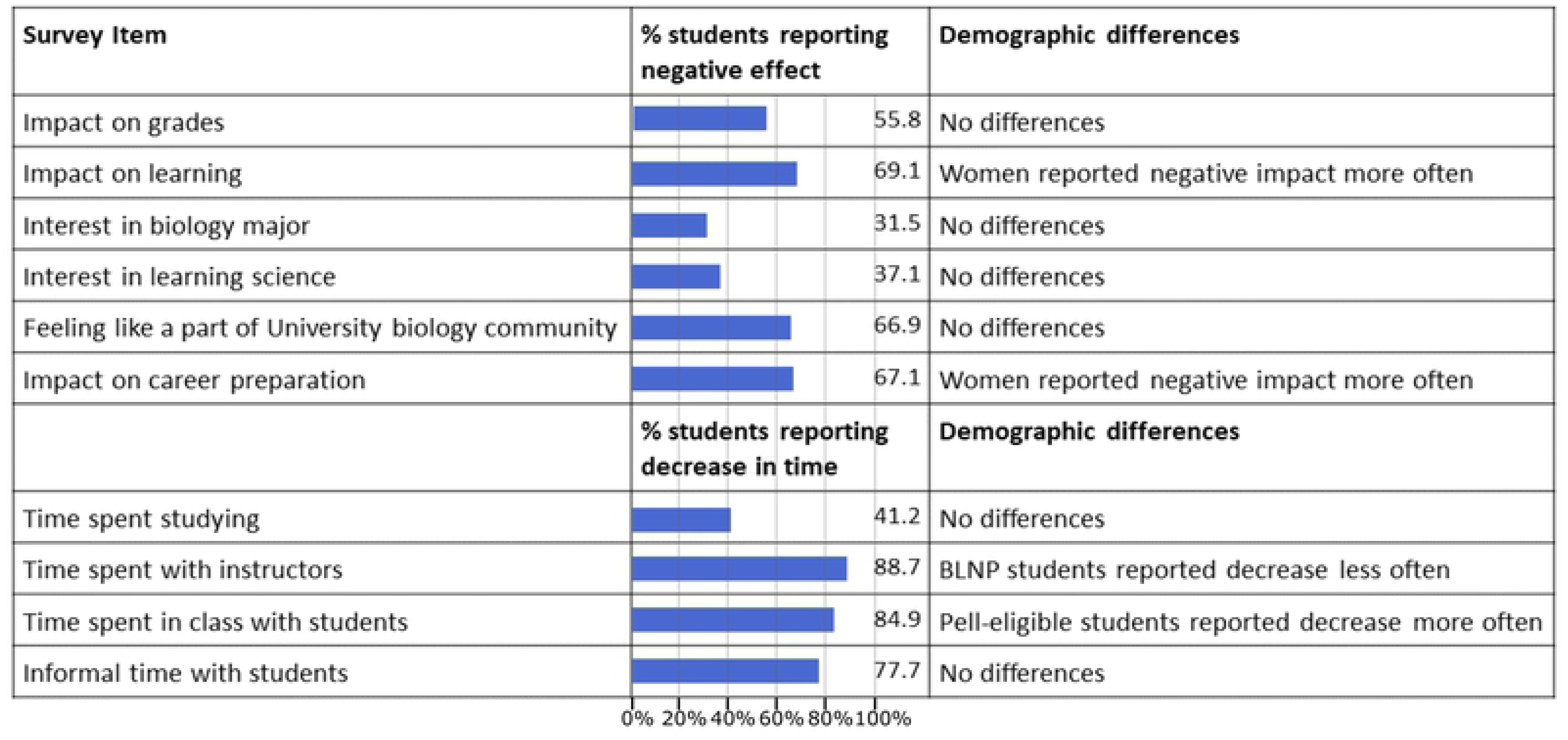
Percentage of students who reported a negative impact or reported a decrease in the time spent on various activities during Spring 2020 along with ordinal regression results on demographic differences. BLNP refers to Black, Latinx, Native American, and Pacific Islanders. Pell eligibility and college generation status were included as proxies for socioeconomic status. The reference groups for the regression analyses were: men, non-BLNP students, continuing-generation students and students that were not eligible for Pell grants.

Our analyses of the closed-ended Likert scale data showed that a large proportion of students (67.1%) reported that the transition to remote learning in Spring 2020 had a negative impact on their career preparation. A relatively smaller but still significant proportion of students reported a negative impact of the transition to remote learning on their interest in their biology major (31.5%) or interest in learning about scientific topics (37.1%). However, many more students (66.9%) reported a negative impact of the transition to remote learning on their feeling of being a part of the biology community at the university. See the Supplemental Materials for full Likert scale response for each of the survey items.

Most students reported that the amount of time they spent on interactions with instructors and other students, both in and outside of class, decreased as a result of the transition to remote learning. In fact, about 63% of students said that the amount of time they spent interacting with other students in class and outside of class greatly decreased in Spring 2020, which was the strongest response option (Table S11). However, student responses were fairly split on the amount of time spent studying for a course, with about 45% of students reporting an increase in the amount of time they spent studying and 41% reporting a decrease (Fig 1).

We collected student and instructor data on instructional practices for eight courses. Among these, only four of the eight instructors agreed that they took steps to reduce cheating in their course and the percentage of students in a given course taught by one of these four instructors who agreed that their instructor took some steps to reduce cheating ranged from 90 to 100%. However, even for the courses where instructors disagreed that they took steps to reduce cheating, 83 to 86% of students agreed that their instructor took some steps to reduce cheating (Figure S1). By contrast, all eight of these instructors agreed that they tried to make their course more flexible. However, there was more variation in student response to whether their instructor tried to make the course more flexible with the percentage of students who agreed with this statement ranging from 61 to 91% across the eight courses. All but one instructor agreed that they tried to make the course easier, but the student agreement with this question was again mixed ranging from 47 to 81% across the courses. Overall, these data show that students tended to slightly overestimate instructor efforts to reduce cheating and slightly underestimate instructor efforts to make the course easier and more flexible.

We did not find significant demographic differences in the student Likert responses to most of the survey items. In Fig 1, we describe the few demographic differences we found through our ordinal mixed models (see full ordinal regression results in the Supplemental Materials). Although most students reported that the time spent with instructors decreased or greatly decreased during the pandemic, the proportion of BLNP students that chose these options was lower than non-BLNP students. Pell-eligible students were more likely to report that time spent with other students in class greatly decreased compared to students that were not Pell-eligible. Lastly, women were significantly more likely than men to report negative impacts on their learning in a course and on career preparation.

## Discussion

Contrary to our predictions, transition to remote learning due to the COVID-19 pandemic in Spring 2020 did not have a negative effect on student grades and instead had a small positive effect across demographic groups among students enrolled in the in-person and online biology degree programs. Our instructor surveys showed that instructors who had to transition to remote learning increased flexibility and made several other changes in assessment practices that might have contributed to the slight increase in student grades in the in-person courses. Despite this increase in grades, our student surveys revealed several negative impacts of the transition to remote learning, particularly on students’ perceived understanding of course content, interactions with other students and instructors, feeling like a part of the biology community at the university, and career preparation. These negative impacts do not seem to have a stronger effect on students with certain social identities over others for the most part. However, women were more likely to report negative impacts on their learning and career preparation compared to men, a result consistent with concerns about widening gender inequities due to the COVID-19 pandemic. Additionally, Pell-eligible students reported a decrease in the amount of time spent in and outside of class interacting with other students more often, which is consistent with concerns regarding logistical difficulties for students from less wealthy backgrounds. Together these findings suggest that instructor responses were effective in mitigating negative impacts on student grades across all demographic groups examined in this study, and notably did not seem to induce any new inequities based on demographics, but that the abrupt transition to remote learning still led to a diminished perception of learning and career development during the Spring 2020 semester for many students.

The observed mismatch between grades and student perceptions of their learning might be because students underestimated their learning [62]. Some studies have shown that student perceptions of learning can be positively correlated with their grades [63–65]. However, a recent study comparing the effects of active and passive (i.e., lectures) instruction on student learning found that students who received active instruction scored higher on the learning assessment but perceived that they learned less than their peers who received passive instruction [66]. Thus, even though it has been shown that students, on average, learn more from active learning [67,68], students’ perception of learning might not match their actual learning. A meta-analysis showed that student perceptions of their learning are more strongly related to affective outcomes, such as motivation and satisfaction, and have a much weaker relationship to learning outcomes, such as scores [69]. However, one reason for this may be that grades are often not an accurate measure of student learning [70]. Given this background and our results that instructors were more flexible with grading after the transition to remote learning in Spring 2020, we think it is likely that the increase in grades does not actually reflect an increase in student understanding of the course material. In contrast, students earned higher grades while self-reporting that they learned less, which we find concerning for the extent to which their completion of these college courses is preparing them for their future careers.

The slight increase in average student grades in Spring 2020 compared to previous semesters is consistent with other studies that have examined student grades in Spring 2020 at other institutions [45–47]. Interestingly, this increase in student grades was observed both in courses that experienced the emergency transition to remote learning and courses in the online degree program that did not experience a transition in modality. Although we did not survey the online instructors, this suggests that both in-person and online instructors may have been responsive to the public health and economic crisis due to the COVID-19 pandemic and became more lenient and flexible in their grading. The increase in student grades was seen across all demographic groups. More specifically, women, Black, Latinx, Native American, Multiracial and Asian students, and Pell-eligible students experienced a similar or slightly larger positive shift in grades as men, white students, and students who were not eligible for Pell grants. Thus, the grade increase in Spring 2020 did not fall along the lines of power and privilege in our society and benefited students with all social identities. A similar result was found in a study on student scores at Victoria University in Australia where the researchers found statistically significant but very small differences in the impact of COVID-19 on student scores between demographic groups [45].

The instructor surveys show that among our study population, most instructors made accommodations related to deadlines and stated that they took steps to make their courses easier for students to do well. Other studies have also reported greater flexibility among instructors in Spring 2020, including instructors in general chemistry courses at a liberal arts college in the US [28]. A survey study of faculty members and administrators across the US found that 64% of faculty members changed the kinds of exams or assignments they asked students to complete in the course and about half of them lowered expectations on the amount of work from their students in the Spring 2020 semester [3]. Additionally, many universities expanded access to pass/fail grading structure instead of the more traditional A-F letter grades for students, with some institutions even making the pass/fail grading structure mandatory for all courses [44]. Arizona State University allowed faculty members to use the range of grading options that have always been available but perhaps not used as often prior to Spring 2020. That included the traditional A through E grading scale, plus the use of the I or Incomplete grade (allowing students to complete coursework within 1 year of the end of the term) and the Y grade which indicates “Satisfactory” work at a level of C or higher, similar to the Pass grade at other universities. Thus, our study affirms other reports that the focus across colleges and universities to make courses more flexible and less stressful for students in Spring 2020 may have off-set potential drops in student grades. While we see the benefit of this flexibility for students, particularly that we did not see demographic differences in these grade increases, we do find it concerning that students still felt as though they learned less. We encourage instructors to be thoughtful of what they are doing to make their courses flexible while maintaining the quality of teaching and providing students with ways to engage in deep learning so that they are not disadvantaged at a later timepoint because they have not learned as much as they needed to in that earlier course.

Many students recognized the positive impact of greater instructor flexibility and changes in assessment practices on their grades, while recognizing the negative impact of the transition on their understanding of the course material. This is consistent with other survey studies that show that students perceived a negative impact on their learning or were less satisfied with their learning after the transition to remote learning [25,41,45]. In our study, most students also reported negative impacts on interactions with other students and instructors, career preparation, and a feeling of being a part of the biology community at the university. These are also consistent with other studies on student experiences [25,43]. A larger survey study of in-person students at Arizona State University across various degree programs, the same institution where our study was conducted, found several striking negative impacts on career preparation due to COVID-19. According to this study, 13% of students delayed graduation, 40% suffered the loss of a job/internship, and 29% of students expected to earn less by age 35 [48].

We found similar perceptions of negative impacts on student learning, interactions and career preparation across demographic groups with few significant differences. We found that women were more likely to report negative impacts on their learning and career preparation compared to men. This is not surprising given the greater childcare obligations with school closures and that women spend more time doing unpaid care work compared to men [32]. In an interview study of engineering students, women reported having to spend more time on domestic duties while men described having more free time after the transition to remote learning during the Spring 2020 semester [71]. Together this suggests that the COVID-19 pandemic has exacerbated gender inequities and could have long-term negative impacts on women’s education and careers that are not captured in simply examining student course grades. We encourage future studies to explore how the COVID-19 pandemic affected the persistence of women in STEM careers.

The only survey item in which we found a significant difference between BLNP and non-BLNP students was the time spent with instructors, where BLNP students chose the option “greatly decreased” less often. Previous studies show that BLNP students often have more interactions with faculty members compared to white students, although they also have negative interactions with faculty members more often [72,73]. Still, their greater experience of interacting with faculty members might have prepared them better to communicate with instructors during emergency remote learning. High-quality interactions with faculty members have been shown to have positive effects on student learning [72,74,75]. However, BLNP students did not report less negative impacts on learning compared to non-BLNP students. This suggests that even though BLNP students reported a decrease in the time spent with instructors less often, it might not have translated into benefits for their learning.

We also found that students from less wealthy backgrounds (operationalized through federal Pell grant eligibility) more often reported a reduction in time spent with other students in class after the transition to remote learning. Pell-eligible students were also 1.2 times more likely to be working a job after the transition to remote learning and 1.5 times more likely to be working more than 20 hours a week compared to students that were not eligible for federal Pell grants (Table S12). With greater availability of recorded lectures, Pell-eligible students may have attended fewer synchronous sessions, thus further reducing their interactions with other students. Although the decrease in interactions with other students is not desirable, making lectures available for students to watch later might offer students greater flexibility in juggling coursework with other work/family responsibilities. Indeed, some students reported positive impacts on learning after the transition to remote learning due to the availability of recorded lectures and being able to learn at their own pace (Table 6). Overall, instructors may need to find a balance between asynchronous learning to make learning more accessible with synchronous learning to foster peer interactions.

**Table 6.**
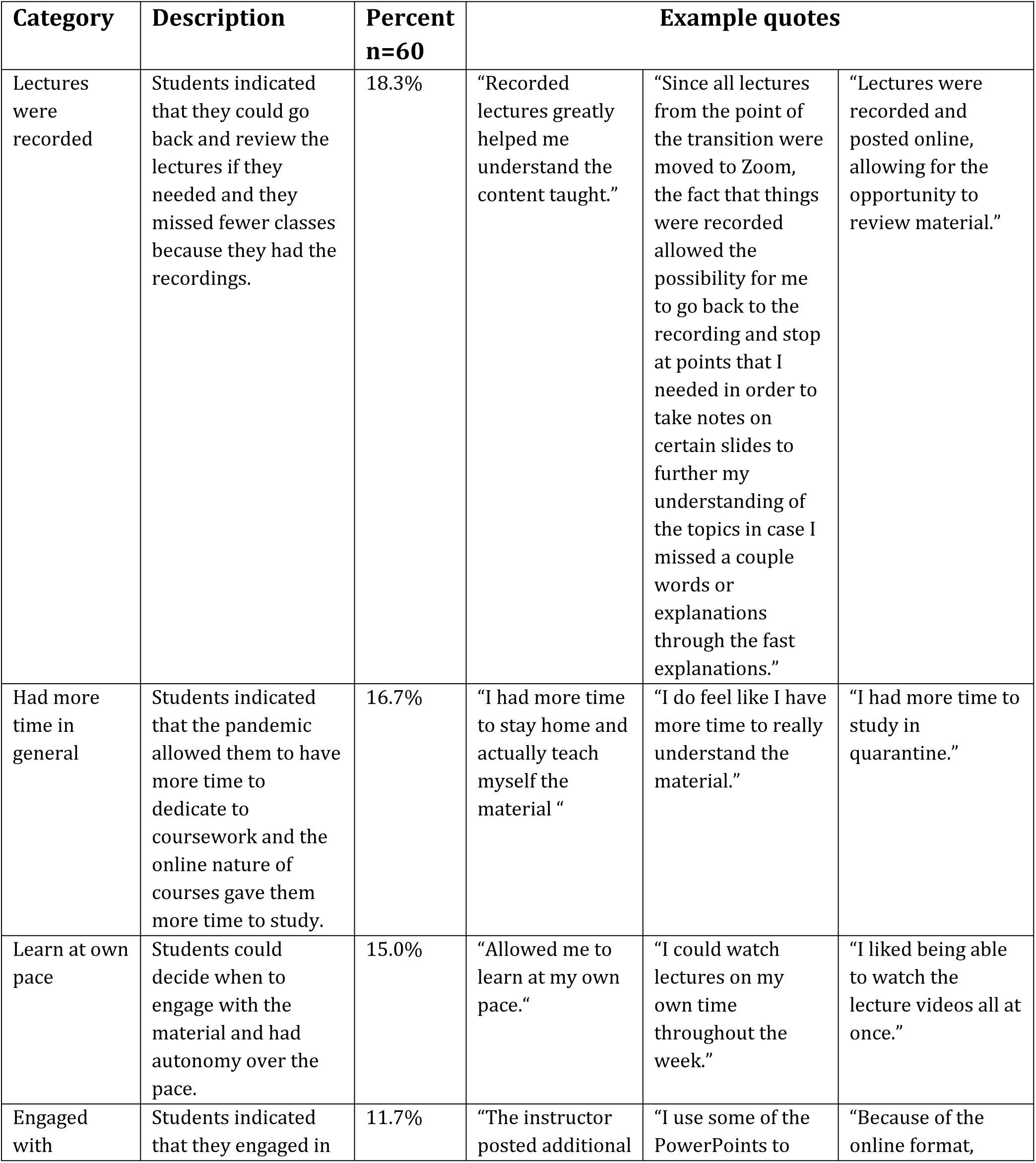
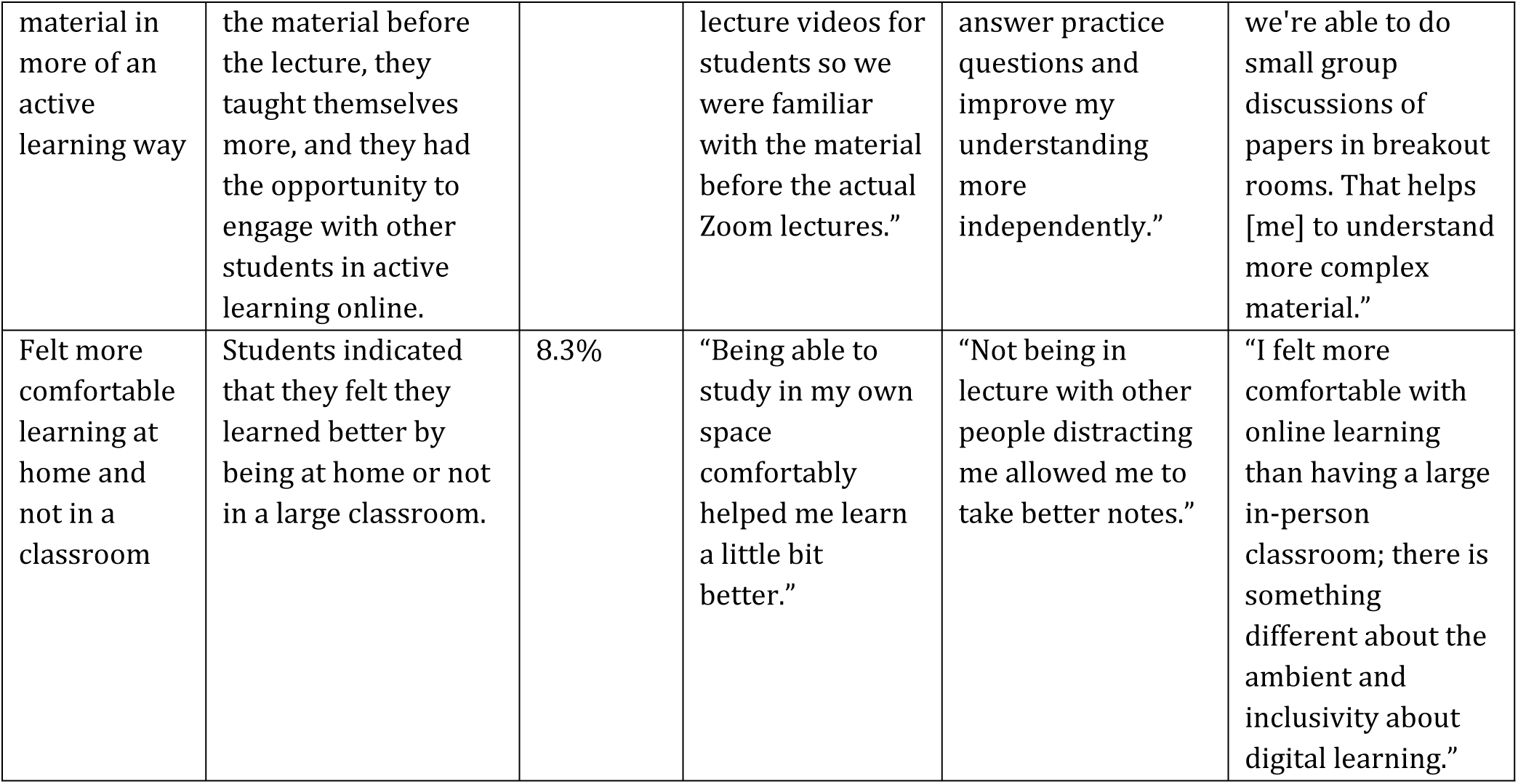
Positive impacts of the transition to remote learning on student learning experiences.

**Table 7.**
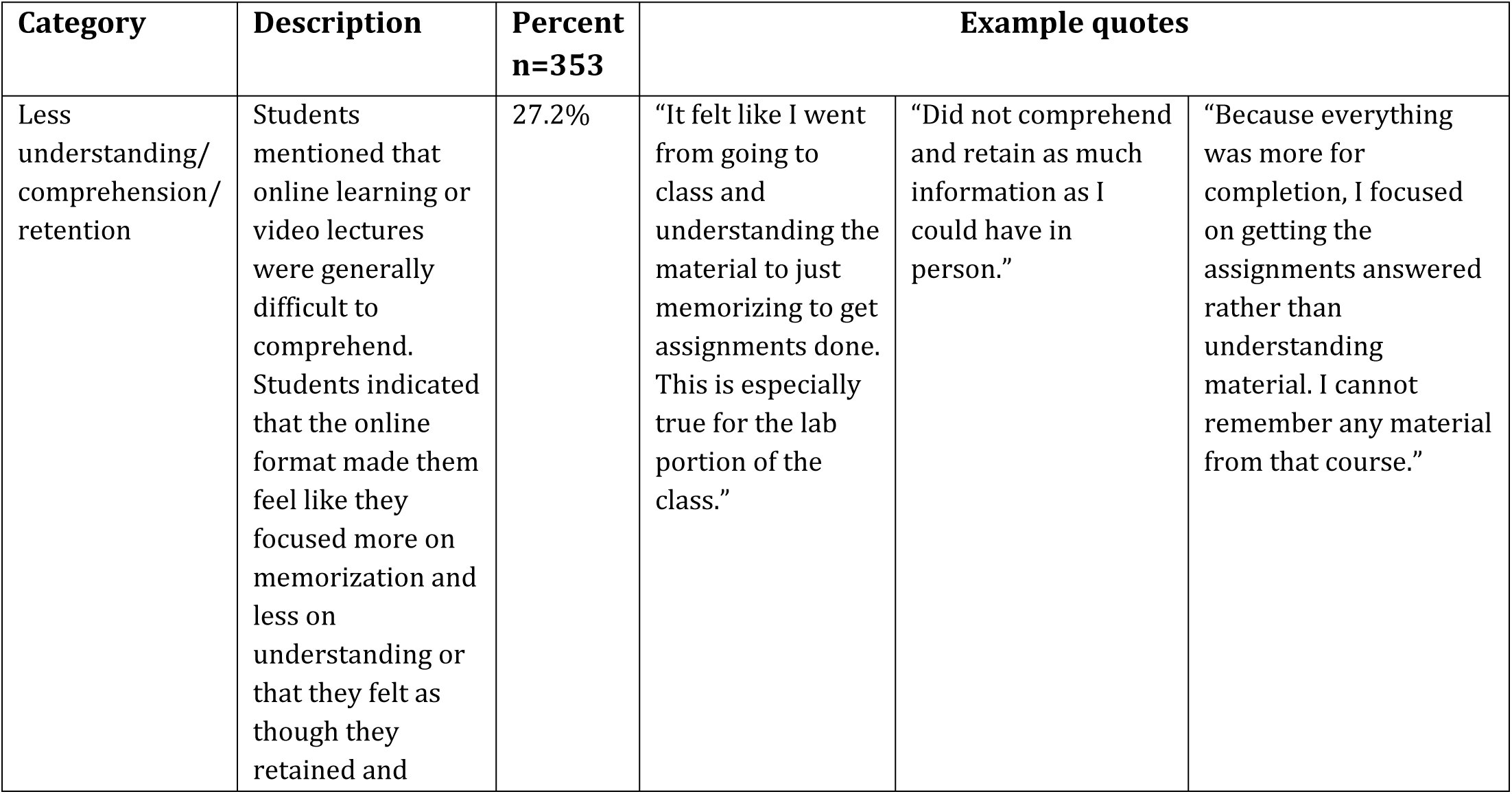
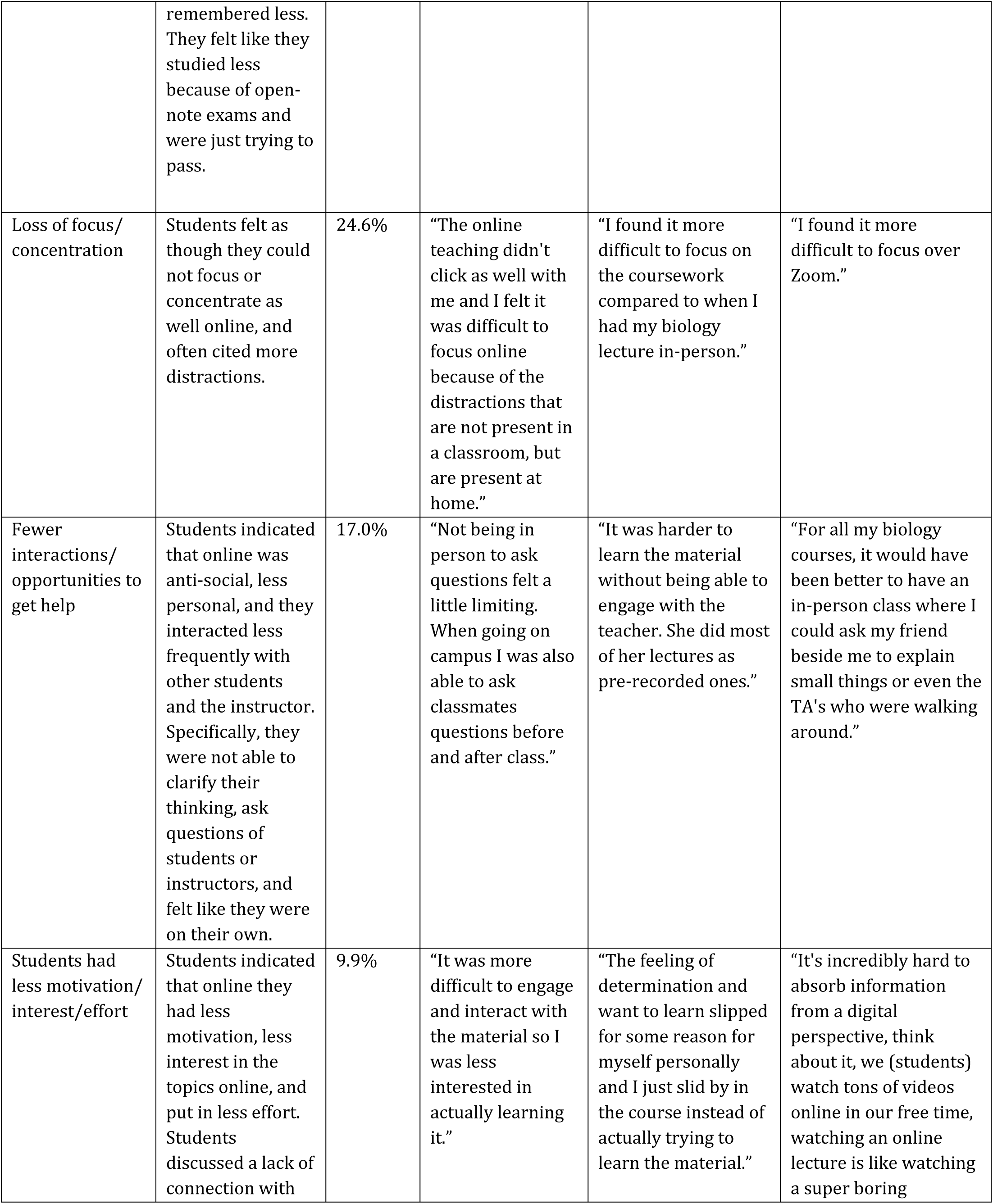
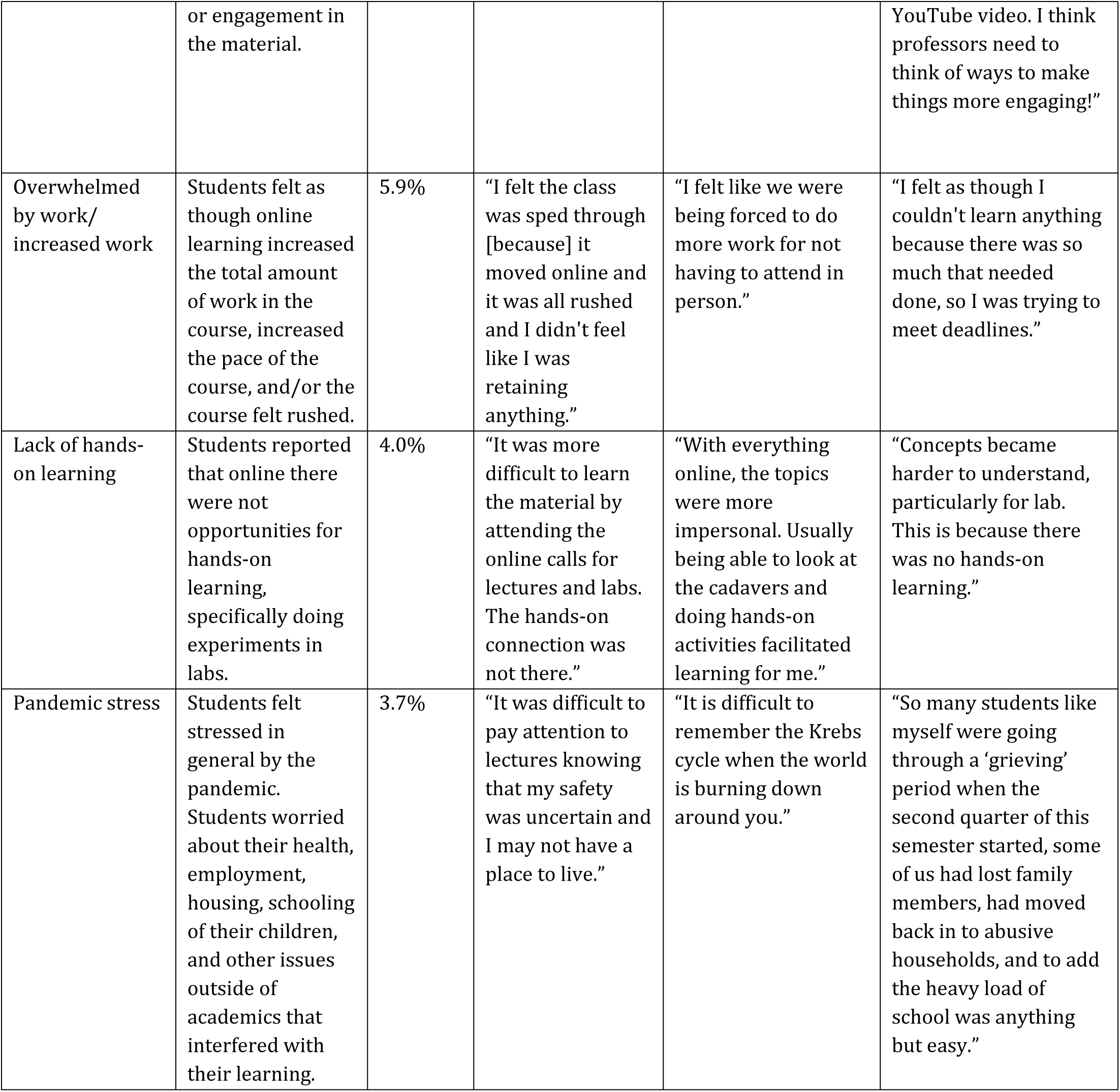
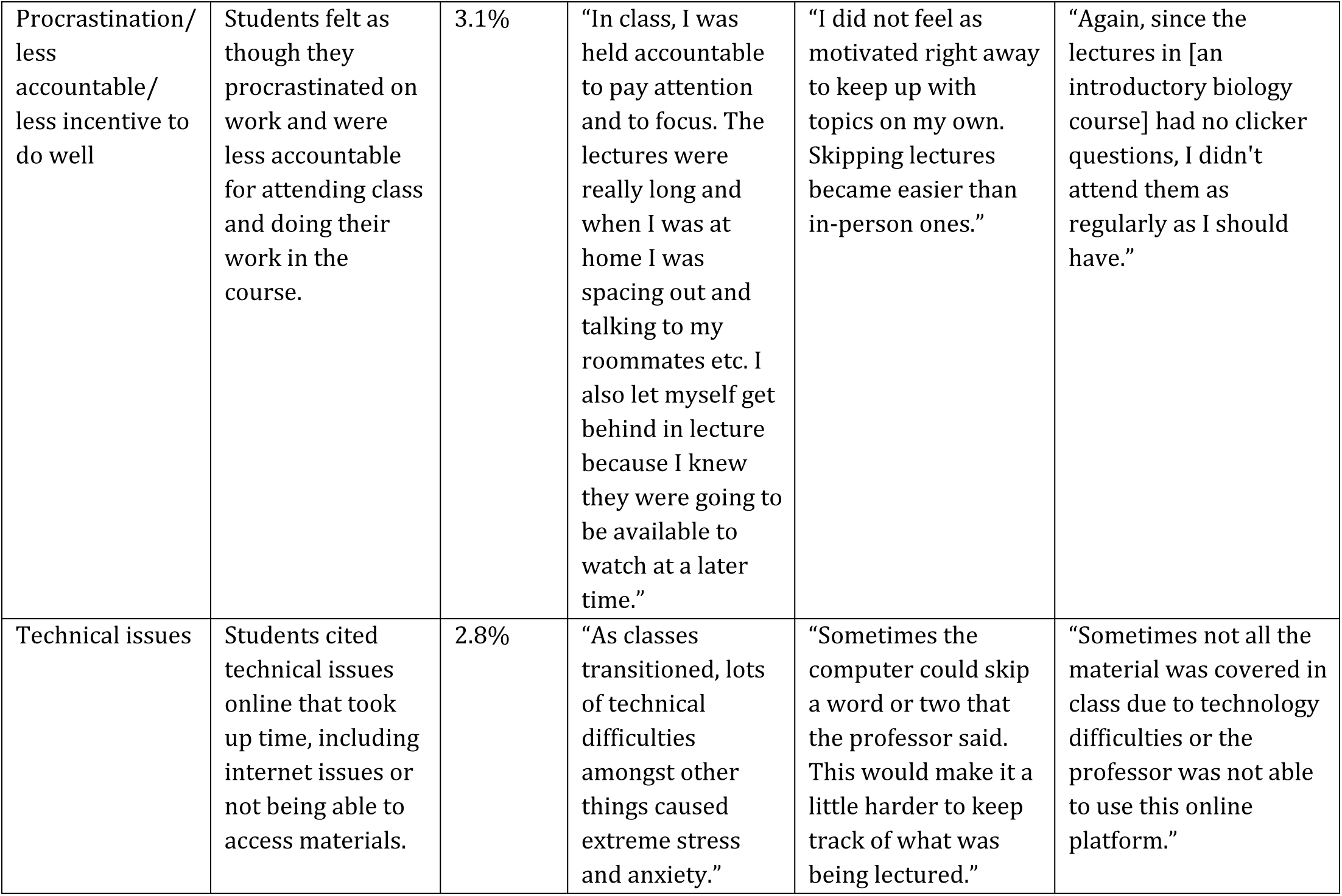
Negative impacts of the transition to remote learning on student learning experiences.

The transition to remote learning had a negative impact on students’ interest in their biology major or interest in learning about scientific topics in about a third of the students. A similar study of students enrolled in a general chemistry course at a large public university in the southern United States found no significant change to students’ identities and intention to pursue a career in science due to COVID-19 [76]. However, we did not find any demographic differences in student responses to questions about science interest, which is encouraging given the importance of increasing representation of women, Black students, Latinx students, and students that grew up in low-income households in STEM. Almost two-thirds of students reported a negative impact of the transition to remote learning on students’ feelings of being a part of the biology community at the university, which is alarming, although not surprising, given that students reported spending less time interacting with both instructors and their peers.

Creating opportunities for increasing interactions using various modes of synchronous and asynchronous communication (e.g., online office hours, discussion boards, apps) might help students feel a greater sense of community and social presence of others in the class.

Instructor responses to our survey items about whether they took steps to prevent cheating, increased flexibility, or made the course easier are in broad agreement with student responses to those survey items. Most students seem to recognize their instructors’ efforts during the transition to adapt their courses to the online modality as well as the public health and economic crisis. However, students’ underestimation of instructor flexibility and changes to make courses easier suggests that communication between students and instructors might need to be strengthened. Instructors may have needed to use more “instructor talk,” which is defined as any discussion that is not specific to the course content, to signify the changes that they were making to the courses and why they were making these changes [77]. It is also possible that the steps that instructors took might not have been sufficient to reach students’ needs or expectations. Because instructors tend to be in better financial situations than their students, perhaps they underestimated some of the student challenges. Setting up robust systems of communication among students, instructors, student support staff members, and administrators might improve the academic climate for all stakeholders and prepare us better for future emergencies or needs to change instruction rapidly. Indeed, an interview study with engineering students found that faculty members communicating care and increasing flexibility was a key element for supporting students [71]. In another study, students indicated the need for constant communication from instructors during remote learning [78]. Thus, developing stronger communication with students and improving “instructor social presence” in online courses, i.e., the sense that the instructor is connected and available for interactions is critical [79–81]. This may be done through casual conversations in discussion boards, leveraging social media and using time in class and during office hours to build classroom community.

## Limitations

Prior work shows that grades are not an accurate measure of student learning, thus we are limited in our ability to accurately measure the effects of the abrupt transition to remote learning due to COVID-19 on student learning [70,82,83]. Moreover, student perceptions of negative impacts of the transition on their learning that we observed might be attributed to the abrupt transition itself or the difficulty of learning during a pandemic. Surveying students in the online program about their experiences in the Spring 2020 semester could have helped us tease apart these two factors more.

Another limitation of our study is the relatively small sample size for our survey dataset which caused us to group data from Black/African American, Hispanic/Latinx, Native American/Alaska Native and Pacific Islander/Native Hawaiian students for analyses. The histories and experiences of racial oppression of these groups in the United States are different from each other and grouping them together erases these differences. Similarly, grouping white and Asian students together into a group is problematic as well, because there are several different ethnicities included in the category of “Asian” in the US which includes ethnicities that are underrepresented in STEM in the US [84]. Despite limited statistical power, we ran ordinal regressions on the survey data with disaggregated race/ethnicity data and have included the results in the Supplemental Materials. We found some significant effects by race/ethnicity in those analyses. Specifically, Asian students perceived being negatively impacted less often on grades, sense of community and career preparation. Also, Black students reported a positive impact on the amount of time studying more often and multiracial students reported a negative impact on grades more often.

Finally, the indicators of socioeconomic status we used (federal Pell grant eligibility and first-generation status) are coarse measures that do not capture socioeconomic status accurately. However, these were the only indicators that we could access from the university registrar.

## Beyond COVID-19: preparing for the next emergency

Instructors responded with greater flexibility in grading in response to the rapid transition to remote learning in Spring 2020 and students received higher grades on average. This shows that instructor response was effective in preventing grade declines for students and doing so equitably across the student population. However, student perceptions of the Spring 2020 semester were less positive, including a sense of diminished learning, loss of community, and reduced career preparation. Even if students’ perception of their learning is not accurate, perceived learning losses might still have important effects on students’ confidence in the course content or interest in pursuing a career in biology. Similar learning losses may have occurred in the Fall 2020 semester and Spring 2021 as the COVID-19 pandemic continued to spread in the US and worldwide.

As we look ahead, these students affected by the pandemic may need more support in subsequent courses, especially in courses that build on prior learning. Dedicating class time to reminding students of important concepts at the beginning of each course or course module could be one form of support. However, upper-level courses may not have class time to spare, so adding supplemental tutorials or instruction may be an alternative way to counteract these potential learning deficits of pre-requisite knowledge. Further, the loss of feeling a part of the biology community needs to be addressed. More intentional community building exercises in classes or in the larger department outside of classes could be ways to heal the damage to students’ sense of belonging.

Although COVID-19 may only affect college education for a particular timeframe, it is important to garner lessons from this experience to prepare for the next emergency, which could be global such as a pandemic, or local such as a natural disaster. Building robust networks of communication among students, instructors, and staff members, and offering greater training and support for online teaching for instructors are steps that could help us prevent some of the challenges associated with the rapid transition to remote learning experienced during the COVID-19 pandemic. We hope that some of the flexibility afforded to students during the pandemic is carried on even after in-person courses resume as instructors may have a better understanding of the myriad of challenges that college students experience daily. Lastly, as the COVID-19 pandemic reminded us, our classrooms and universities do not exist in isolation and are a part of the larger society and therefore, affected by the larger societal forces and power structures that impact student learning in our institutions. Therefore, we must continue to strive toward social justice inside and outside our higher education institutions.

## Acknowledgements

We would like to thank the instructors who filled out our survey and those who distributed our survey to students. We would also like to thank the students who filled out our survey and shared their experiences. Lastly, we want to thank the instructors and undergraduate researchers who participated in our think-aloud interviews to establish cognitive validity of our survey instruments.

## Supporting Information

**S1 File. Instructor survey.docx** Instructor survey questions analyzed in this study.

**S2 File. Student survey.docx** Student survey questions analyzed in this study.

**S1 Table.** Number of students in the courses analyzed for the grades data.

**S2 Table. Regression model specifications.** Model 1 estimates the Spring 2020 effect and its interaction with instruction mode. Model 2 extends Model 1 to consider the possibility of demographic interaction effects. Model 3 estimates the predictive value of instructors’ reported course changes on the Spring 2020 grade shift as compared to the same course in Spring 2018/19.

**S3 Table. Linear regression results for courses in in-person and online degree program.** This model shows interaction effects among each demographic category and the COVID-19 semester.

**S4 Table. Changes in assessment practices.** Frequency at which instructors chose various options regarding changes made to their assessment practices following the transition to remote learning in Spring 2020.

**S5 Table. Ordinal regression results for the perceived impact of the transition to remote learning on students.** Ordinal regression output with survey items about impact on students as the outcomes and demographics as predictors.

**S6 Table. Ordinal regression results for the perceived impact of the transition to remote learning on student interactions.** Ordinal regression output with survey items about impact on time spent studying and interactions as the outcomes and demographics as predictors.

**S7 Table. Ordinal regression results for perceived impact on students with disaggregated race/ethnicity data.** Ordinal regression output with survey items about impact on students as the outcomes and demographics as predictors with disaggregated race/ethnicity data.

**S8 Table. Ordinal regression results for perceived impact on student interactions with disaggregated race/ethnicity data.** Ordinal regression output with survey items about impact on time spent studying and interactions as the outcomes and demographics as predictors with disaggregated race/ethnicity data.

**S9 Table. Impact on grades, learning and career preparation.** Full distribution of Likert scale responses on the student survey on the perceived impact of the transition to remote learning in Spring 2020 on grades, learning and career preparation.

**S10 Table. Impact on interest in biology major, interest in learning science and sense of community.** Full distribution of Likert scale responses on the student survey on the perceived impact of the transition to remote learning in Spring 2020 on interest in biology major, interest in learning science and sense of community.

**S11 Table. Impact on time spent studying and interacting with others.** Full distribution of Likert scale responses on the student survey on the perceived impact of the transition to remote learning in Spring 2020 on time spent studying and interacting with others.

**S12 Table. Pell-eligible and students who worked a job in the student survey dataset.**

**S1 Figure. Distribution of student responses about instructional practices.** Blue points indicate the option that the instructor for the course chose.

**S2 Figure. Distribution of student responses on perceived impact of transition to remote learning on grades by social identities.** Bar plot showing student responses by gender, race/ethnicity and socioeconomic status on the perceived impact of the transition to remote learning in Spring 2020 on their grades. BLNP refers to Black, Latinx, Native American, and Pacific Islanders. Pell eligibility and college generation status are included as proxies for socioeconomic status.

**S3 Figure. Distribution of student responses on perceived impact of transition to remote learning on learning by social identities.** Bar plot showing student responses by gender, race/ethnicity and socioeconomic status on the perceived impact of the transition to remote learning in Spring 2020 on their learning. BLNP refers to Black, Latinx, Native American, and Pacific Islanders. Pell eligibility and college generation status are included as proxies for socioeconomic status.

**S4 Figure. Distribution of student responses on perceived impact of transition to remote learning on interest in biology major by social identities.** Bar plot showing student responses by gender, race/ethnicity and socioeconomic status on the perceived impact of the transition to remote learning in Spring 2020 on their interest in biology major. BLNP refers to Black, Latinx, Native American, and Pacific Islanders. Pell eligibility and college generation status are included as proxies for socioeconomic status.

**S5 Figure. Distribution of student responses on perceived impact of transition to remote learning on interest in learning about scientific topics by social identities.** Bar plot showing student responses by gender, race/ethnicity and socioeconomic status on the perceived impact of the transition to remote learning in Spring 2020 on their interest in learning about scientific topics. BLNP refers to Black, Latinx, Native American, and Pacific Islanders. Pell eligibility and college generation status are included as proxies for socioeconomic status.

**S6 Figure. Distribution of student responses on perceived impact of transition to remote learning on feeling a part of the biology community at the university by social identities.** Bar plot showing student responses by gender, race/ethnicity and socioeconomic status on the perceived impact of the transition to remote learning in Spring 2020 on their feeling a part of the biology community at the university. BLNP refers to Black, Latinx, Native American, and Pacific Islanders. Pell eligibility and college generation status are included as proxies for socioeconomic status.

**S7 Figure. Distribution of student responses on perceived impact of transition to remote learning on career preparation by social identities.** Bar plot showing student responses by gender, race/ethnicity and socioeconomic status on the perceived impact of the transition to remote learning in Spring 2020 on their career preparation. BLNP refers to Black, Latinx, Native American, and Pacific Islanders. Pell eligibility and college generation status are included as proxies for socioeconomic status.

**S8 Figure. Distribution of student responses on perceived impact of transition to remote learning on amount of time spent studying by social identities.** Bar plot showing student responses by gender, race/ethnicity and socioeconomic status on the perceived impact of the transition to remote learning in Spring 2020 on their amount of time spent studying. BLNP refers to Black, Latinx, Native American, and Pacific Islanders. Pell eligibility and college generation status are included as proxies for socioeconomic status.

**S9 Figure. Distribution of student responses on perceived impact of transition to remote learning on amount of time spent interacting with instructors by social identities.** Bar plot showing student responses by gender, race/ethnicity and socioeconomic status on the perceived impact of the transition to remote learning in Spring 2020 on their amount of time spent interacting with instructors. BLNP refers to Black, Latinx, Native American, and Pacific Islanders. Pell eligibility and college generation status are included as proxies for socioeconomic status.

**S10 Figure. Distribution of student responses on perceived impact of transition to remote learning on amount of time interacting with other students in class by social identities.** Bar plot showing student responses by gender, race/ethnicity and socioeconomic status on the perceived impact of the transition to remote learning in Spring 2020 on their amount of time spent interacting with other students in class. BLNP refers to Black, Latinx, Native American, and Pacific Islanders. Pell eligibility and college generation status are included as proxies for socioeconomic status.

**S11 Figure. Distribution of student responses on perceived impact of transition to remote learning on amount of time interacting with other students outside of class by social identities.** Bar plot showing student responses by gender, race/ethnicity and socioeconomic status on the perceived impact of the transition to remote learning in Spring 2020 on their amount of time spent interacting with other students outside of class. BLNP refers to Black, Latinx, Native American, and Pacific Islanders. Pell eligibility and college generation status are included as proxies for socioeconomic status.

